# SPOT-1D-LM: Reaching Alignment-profile-based Accuracy in Predicting Protein Secondary and Tertiary Structural Properties without Alignment

**DOI:** 10.1101/2021.10.16.464622

**Authors:** Jaspreet Singh, Kuldip Paliwal, Jaswinder Singh, Yaoqi Zhou

**Affiliations:** Signal Processing Laboratory, School of Engineering and Built Environment, Griffith University, Brisbane, QLD 4111, Australia; Institute for Glycomics, Griffith University, Parklands Dr. Southport, QLD 4222, Australia; Institute for Systems and Physical Biology, Shenzhen Bay Laboratory, Shenzhen 518055, China; Peking University Shenzhen Graduate School, Shenzhen, 518055, P.R. China

## Abstract

Protein language models have emerged as an alternative to multiple sequence alignment for enriching sequence information and improving downstream prediction tasks such as biophysical, structural, and functional properties. Here we show that a combination of traditional one-hot encoding with the embeddings from two different language models (ProtTrans and ESM-1b) allows a leap in accuracy over single-sequence based techniques in predicting protein 1D secondary and tertiary structural properties, including backbone torsion angles, solvent accessibility and contact numbers. This large improvement leads to an accuracy comparable to or better than the current state-of-the-art techniques for predicting these 1D structural properties based on sequence profiles generated from multiple sequence alignments. The high-accuracy prediction in both secondary and tertiary structural properties indicates that it is possible to make highly accurate prediction of protein structures without homologous sequences, the remaining obstacle in the post AlphaFold2 era.

## Introduction

Recently, Alphafold2 has achieved what was thought impossible: predicted protein structures at experimental accuracy for the majority of target proteins in critical assessment of structure prediction techniques (CASP14)^1^. This revolution was built on accumulating improvement in predicting backbone secondary structure^2–6^ and residue-residue contact maps^7–9^. This success, however, does not mean the protein structure prediction problem is solved, as AlphaFold2 requires a minimum of 30 effective homologous sequences to achieve an accurate structure prediction^1^ and a large portion of proteins lacks homologous sequences^10^. Moreover, sequence-homology search requires increasingly intensive computing time. Thus, it is essential to develop accurate structure prediction methods without relying on homologous sequences. To do this, the first step is to develop accurate alignment-free prediction of protein backbone and other 1D-structural properties with a single sequence as input.

To date, only a few single-sequence-based methods have been developed for protein secondary structure prediction. Examples are PSIPRED-Single^11^, SPIDER3-Single^12^, ProteinUnet^13^, NetSurfP-2.0^4^, and SPOT-1D-Single^14^. PSIPRED-Single predicts the secondary structure only while SPIDER3-Single, ProteinUnet, and SPOT-1D-Single predicts secondary structure, Accessible Surface Area (ASA)^15^, Half-Sphere Exposure (HSE)^16^ and Backbone torsion angles (*ψ, ϕ, θ*, and *τ*). SPIDER3-Single employed iterative learning on a two-layer Bidirectional Long-Short-Term-Memory(LSTM) cells^17^ on a training set of approximately 10000 proteins. ProteinUnet followed the same strategy except replacing the Bi-LSTM model with a convolution-based Unet architecture^18^, which achieved a similar performance but with a smaller computational requirement. More recently, SPOT-1D-Single improved over all previous predictors by taking advantage of a high sequence identity training set and an ensemble of Convolution and LSTM based hybrid to improve the performance on completely independent test sets. Although these single-sequence models do improve over profile-based methods for proteins with a low number of effective homologous sequences (Neff), there is above 10% gap for those sequences with higher Neff values: 74% for three-state secondary structure prediction, compared to 86% by profile-based techniques^14^.

Recently, unsupervised deep learning methods inspired by Natural Language Processing were introduced to extract features from protein sequences^19–22^. These methods were trained on extensive protein databases such as Uniref^23^, Uniclust^24^, Pfam^25^, and BFD^26,27^. One state-of-the-art protein language model (LM) is ProtTrans^22^ trained on the Uniref50 dataset. It employs a transformer-based auto-encoder model T5 to generate the embedding. Another protein language model also trained on Uniref50 dataset is ESM-1b which uses a 34 Transformer model^21^.

In this work, we explored the combined use of ProtTrans and ESM-1b generated embedding to train a downstream predictor of secondary structure and 1D structural properties. We demonstrated that the new alignment-free model can match or exceed the performance of sequence-profile-based prediction of 1D structural properties for both high and low Neff proteins without searching for homologous sequences.

## Results

### Feature Analysis

Our model was built on a combination of three main input features: single-sequence one-hot, ESM-1b, and ProtTrans encodings. We trained three individual neural network models (Two-Layer LSTM, MS-ResNet, and MS-Res-LSTM) with different combinations of these three input features. Results are shown in Figure 1 for three-state (SS3) secondary structure prediction on independent test sets of TEST2018, TEST2020, and Neff1-2020 datasets. TEST2018 (deposited between January 2018 - June 2018) is a set based on the commonly used criterion of <25% sequence identity cutoff from all proteins released before 2018 on the PDB. TEST2020 (deposited between year 2018-2020) is a new harder test set with remote homology removed by HMM-based search (see Methods), whereas Neff1-2020 contains the proteins in TEST2020 with no homologs (Neff=1, 46 proteins). The accuracy on the easy TEST2018 (86.5% by three features) is significantly higher than on the hard set TEST2020 (80%) as expected. At the single feature level, both ESM-1b and ProtTrans encodings are significantly better in predicting secondary structure. ProtTrans has comparable performance to ESM-1b in TEST2018 but a better performance in the more difficult TEST2020 and NEFF1-2020. Adding one-hot encoding to ProtTrans makes marginal improvement over ProtTrans alone on TEST2018 but a larger improvement in TEST2020 and Neff1-2020. On the other hand, adding one-hot encoding to ESM-1b makes a comparable performance on TEST2018 but a worse performance in TEST2020. This surprising result is not observed for eight-state (SS8) prediction (Supplementary Figure S1). What is the most important is that combining three features make a consistent improvement in all three datasets and three networks. The improvement is the largest for the difficult case (TEST2020 and NEFF1-2020). Overall, the three-state (SS3) secondary structure accuracy for all test sets improves 6-10% from 73-76% for a single-sequence-based method (one-hot-encoding) to 79-86% after combining three features. Moreover, the performance on TEST2020 and Neff1-2020 is nearly identical, indicating that unlike profile-based models, the performance of the current alignment-free method is independent of how many homologous sequences a protein has. Similar trends were observed for eight-state (SS8) structure prediction (Supplementary Figure S1), ASA, and HSE (Supplementary Table S1), and backbone torsion angle prediction (Supplementary Table S2).

**Figure 1.**
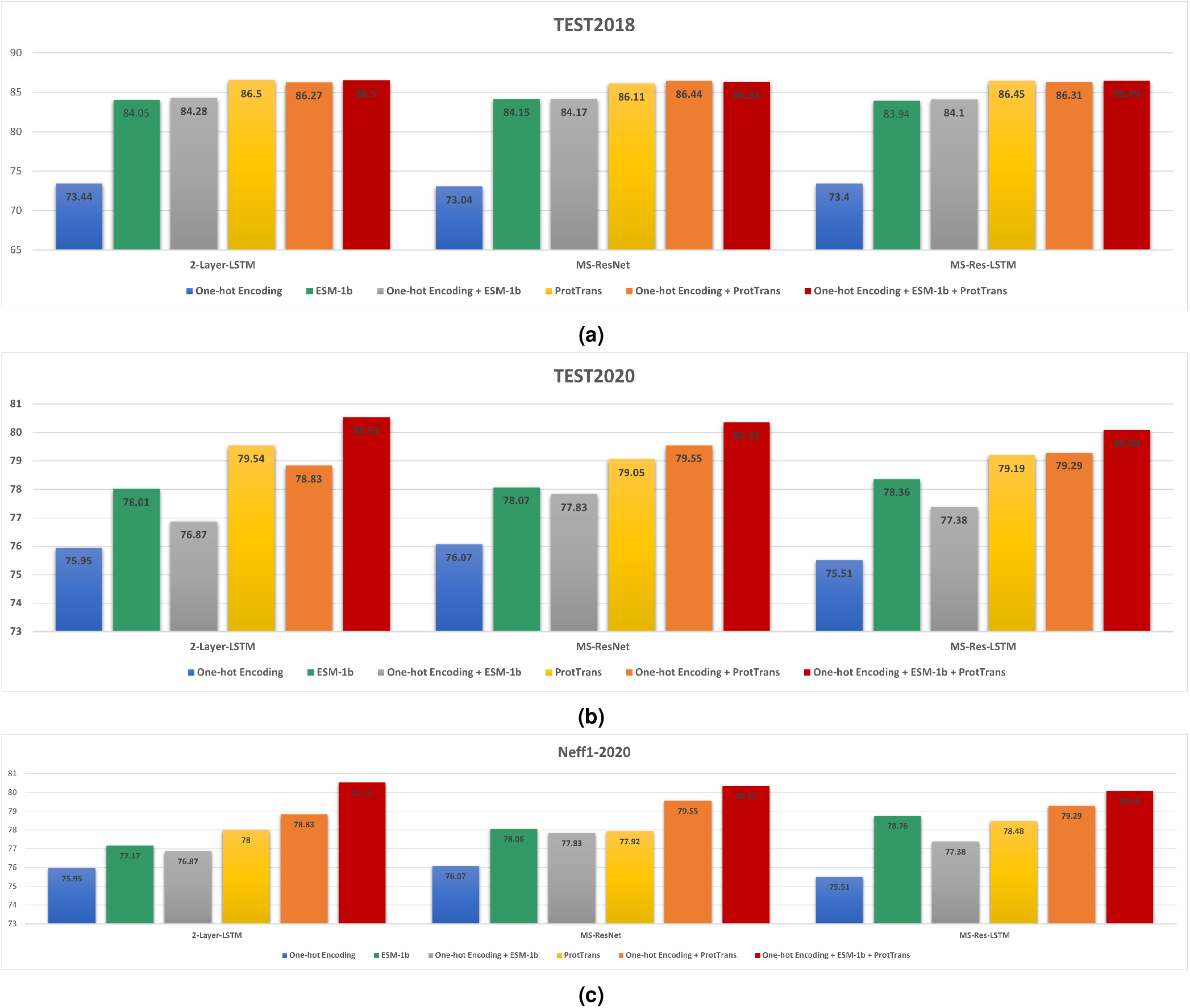
Performance in secondary structure prediction by using different input features as labelled for three different model architectures on three test sets (TEST2018, TEST2020, and Neff1-2020).

### Ensemble learning

The individual models were combined into an ensemble to further improve model performance. To demonstrate the advantage of ensemble learning over individual models, Table 1 presents the results of the selected three models and the results of the ensemble on TEST2018 and TEST2020. As we can see, for all properties tested, the trends we observed in this table is similar to what we observed in our previous work SPOT-1D-Single. The ensemble performance drops the error for *ψ, ϕ, θ*, and *τ* over the best individual model by 0.90%, 0.89%, 1.20%, and 0.87% on TEST2018, respectively. For secondary structure prediction three-state (SS3) and eight-state (SS8) the ensemble accuracy is 86.74% and 76.47%, which is 0.27% and 0.52% better than the best individual model. Similar improvement is also visible in Pearson’s Correlation Coefficient (PCC) for ASA, HSE-U, and CN predictions. The same trend is observed for TEST2020.

**Table 1.** Individual model performance as compared to the ensemble performance on TEST2018 and TEST2020 set for prediction of secondary structure in three (SS3) and eight (SS8) states, solvent accessibility (ASA), half-sphere-exposure-up (HSE-u), half-sphere-exposure-down (HSE-d), contact number (CN), backbone angles (*ψ, ϕ, θ*, and *τ*). Performance measures are accuracy for SS3 and SS8, correlation coefficient for ASA, HSE-u, HSE-d, and CN, and mean absolute errors for the angles.

| Model | TEST2018 |  |  |  |  |  |  |  |  |  | TEST2020 |  |  |  |  |  |  |  |  |  |
| --- | --- | --- | --- | --- | --- | --- | --- | --- | --- | --- | --- | --- | --- | --- | --- | --- | --- | --- | --- | --- |
| | SS3 | SS8 | ASA | HSE-u | HSE-d | CN | $\psi$ | $\phi$ | $\theta$ | $\tau$ | SS3 | SS8 | ASA | HSE-u | HSE-d | CN | $\psi$ | $\phi$ | $\theta$ | $\tau$ |
| 2-Layer-LSTM | 86.50 | 76.07 | 0.804 | 0.745 | 0.755 | 0.788 | 23.964 | 16.139 | 6.540 | 24.821 | 79.57 | 66.44 | 0.708 | 0.516 | 0.591 | 0.612 | 36.792 | 20.671 | 8.738 | 36.149 |
| Multi-Scale ResNet | 86.33 | 75.57 | 0.799 | 0.748 | 0.748 | 0.786 | 24.285 | 16.216 | 6.577 | 25.114 | 79.59 | 66.32 | 0.700 | 0.510 | 0.588 | 0.607 | 37.040 | 20.879 | 8.793 | 36.193 |
| Multi-Scale ResNet LSTM | 86.49 | 75.85 | 0.799 | 0.749 | 0.748 | 0.778 | 24.396 | 16.426 | 6.617 | 25.280 | 79.48 | 66.33 | 0.702 | 0.512 | 0.584 | 0.606 | 36.877 | 20.849 | 8.725 | 36.118 |
| Ensemble (This work) | 86.74 | 76.47 | 0.814 | 0.759 | 0.761 | 0.690 | 23.748 | 15.995 | 6.461 | 24.605 | 79.82 | 66.68 | 0.731 | 0.522 | 0.597 | 0.623 | 36.574 | 20.672 | 8.674 | 35.795 |

### Method comparison

The performance for three-state (SS3) secondary structure prediction given by our ensemble method named SPOT-1D-LM is compared with four single-sequence-based methods PSIPRED-Single, SPIDER3-Single, ProteinUnet and SPOT-1D-Single along with two profile-based methods SPOT-1D and NetSurfP-2.0 on five different test sets (TEST2018, TEST2020, Neff1-2020, CASP12-FM, and CASP13-FM) in Figure 2. The result confirms a large leap from 72-74% by single-sequence-based methods to 80-86% by alignment-profile-based methods for the prediction accuracy for TEST2018, TEST2020, CASP12-FM and CASP13-FM. The performance of profile-based methods is worse than the performance of single-sequence-based methods only for Neff1-2020, confirming the previous finding that profile-based methods lose their accuracy when lacking homologous sequences. Importantly, our language-model-based method achieves a performance that matches or beats those of profile-based methods for all test sets. Furthermore, it improves over single-sequence-based methods even for Neff1-2020. For example, SPOT-1D-LM performs 0.66%, 1.6%, and 17% better than SPOT-1D, NetSurfP-2.0, and SPOT-1D-Single, respectively, for SS3 prediction for TEST2018. Its performance on TEST2020, CASP12-FM and CASP13-FM is comparable to that of the profile-based SPOT-1D and better than that of the profile-based NetSurfP-2.0. Similar trends are also observed for SS8 prediction, as shown in Supplementary Figure S2. The matching performance of SPOT-1D-LM with profile-based models on backbone torsion angles are also illustrated in Table 2, Table 3 for TEST2020, Supplementary Table S3 for CASP12-FM and Supplementary Table S4 for CASP13-FM.

**Table 2.**
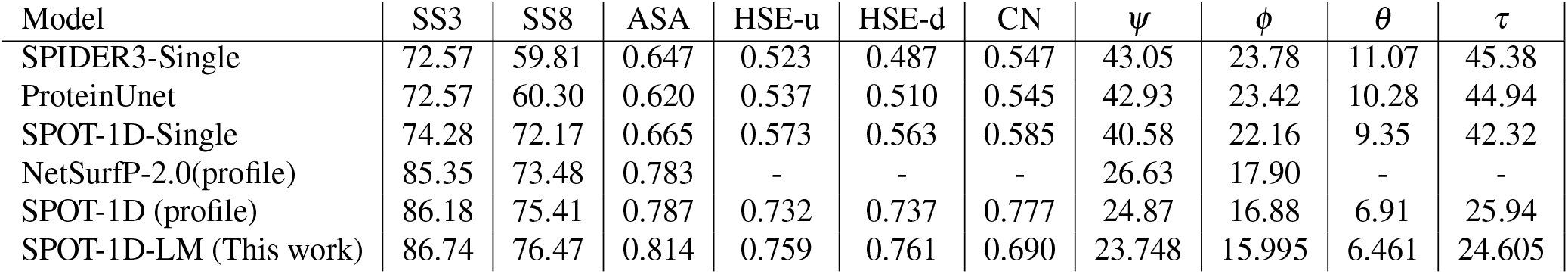
Comparing the performance of SPOT-1D-LM with single-sequence-based methods (SPIDER3-Single, ProteinUnet, and SPOT-1D-Single) and sequence-profile-based methods (SPOT-1D and NetSurfP-2.0) in the prediction of secondary structure in three (SS3) and (SS8) states, solvent accessibility (ASA), half-sphere-exposure-up (HSE-u), HSE-down (HSE-d), contact number (CN), backbone angles(*ψ, ϕ, θ* and *τ*) for TEST2018. Performance measures are accuracy for SS3 and SS8, correlation coefficient for ASA, HSE-u, HSE-d, and CN, and mean absolute errors for the angles.

**Table 3.** Comparing the performance of SPOT-1D-LM with single-sequence-based methods (SPIDER3-Single, ProteinUnet, and SPOT-1D-Single) and sequence-profile-based methods (SPOT-1D and NetSurfP-2.0) in the prediction of secondary structure in three (SS3) and eight (SS8) states, solvent accessibility (ASA), half-sphere-exposure-up (HSE-u), HSE-down (HSE-d), contact number (CN), backbone angles (*ψ, ϕ, θ* and *τ*) for TEST2020. Performance measures are accuracy for SS3 and SS8, correlation coefficient for ASA, HSE-u, HSE-d, and CN, and mean absolute errors for the angles.

| Model | SS3 | SS8 | ASA | HSE-u | HSE-d | CN | $\psi$ | $\phi$ | $\theta$ | $\tau$ |
| --- | --- | --- | --- | --- | --- | --- | --- | --- | --- | --- |
| SPIDER3-Single | 71.31 | 57.57 | 0.596 | 0.358 | 0.417 | 0.434 | 45.64 | 23.48 | 11.52 | 46.04 |
| ProteinUnet | 72.20 | 58.71 | 0.555 | 0.366 | 0.426 | 0.441 | 44.87 | 23.19 | 10.49 | 44.95 |
| SPOT-1D-Single | 73.80 | 60.35 | 0.621 | 0.400 | 0.478 | 0.485 | 44.25 | 22.92 | 9.88 | 43.67 |
| NetSurfP-2.0(profile) | 79.42 | 66.36 | 0.702 | - | - | - | 35.07 | 20.70 | - | - |
| SPOT-1D (profile) | 80.52 | 67.76 | 0.691 | 0.516 | 0.594 | 0.60 | 34.46 | 20.33 | 8.50 | 33.64 |
| SPOT-1D-LM (This work) | 79.82 | 66.68 | 0.731 | 0.522 | 0.597 | 0.704 | 36.57 | 20.67 | 8.67 | 35.80 |

**Figure 2.**
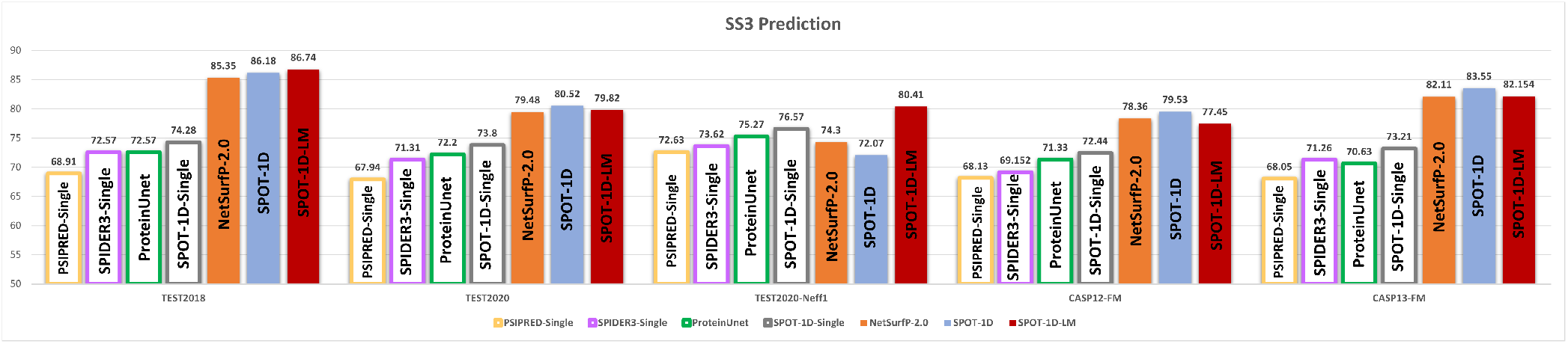
Comparing the accuracy of secondary structure prediction of SPOT-1D-LM (this work) with single sequence methods (SPIDER3-Single, ProteinUnet, and SPOT-1D-Single) and sequence-profile-based methods (SPOT-1D and NetSurfP-2.0) on five test sets (TEST2018, TEST2020, Neff1-2020, CASP12-FM, and CASP14-FM) for three-state (SS3) secondary structure prediction.

Secondary structure is dominated by local interactions. How about structural properties that are based on tertiary structures? Figure 3 examines the performance of different predictors for ASA prediction on five different test sets (TEST2018, TEST2020, Neff1-2020, CASP12-FM, and CASP13-FM). Again, we observe that profile-based methods perform far better than single-sequence-based methods in ASA prediction except when Neff=1 (Neff1-2020). More importantly, SPOT-1D-LM performs the best for all test sets. It outperforms the profile-based method NetSurfP-2.0 by 4%, 4%, 10%, 0.9% and 9% on TEST2018, TEST2020, Neff1-2020, CASP12-FM, and CASP13-FM, respectively. Comparing to SPOT-1D, its improvement is 3%, 6%, 19%, 1%, and 9%, respectively. Better or comparable performance is also observed for other tertiary structural properties such as contact number (CN) and half sphere exposures (HSE-u and HSE-d) as shown in Table 2 for TEST2018, Table 3 for TEST2020, Supplementary Table S3 for CASP12-FM and Supplementary Table S4 for CASP13-FM.

**Figure 3.**
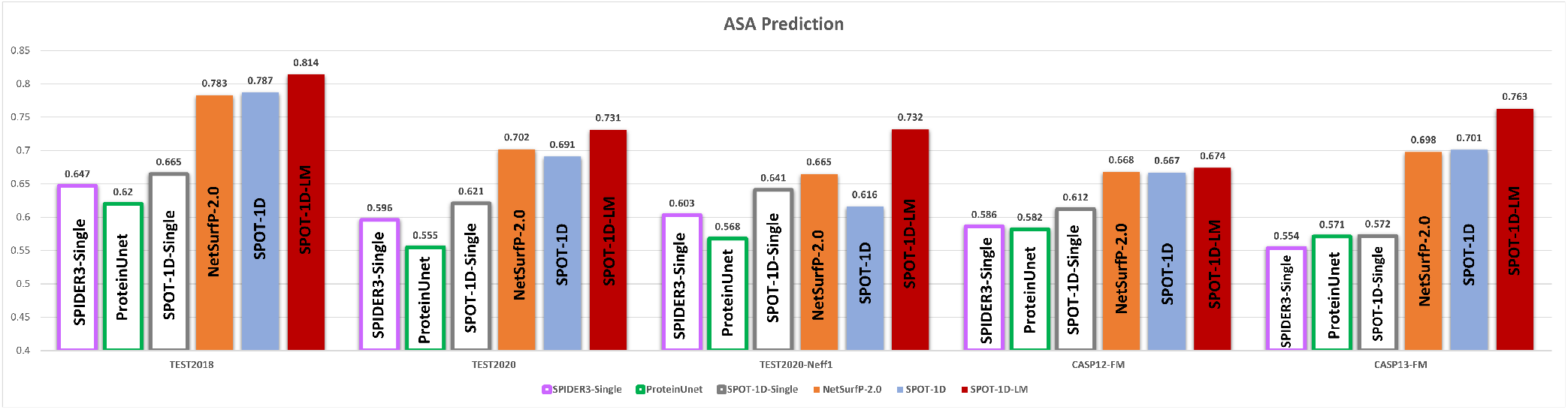
As in Figure 2 but for prediction of tertiary structure proteins (solvent accessibility).

## Discussions

In this paper, we have developed a new Language-model-based method for predicting one-dimensional structural properties of proteins, including secondary structure, solvent accessible surface area, and backbone torsion angles. We employed an ensemble of three network variants of ResNet and LSTM models, trained on approximately 40000 proteins with embedding generated from ESM-1b and ProtTrans. The model is then compared to other predictors on independent and non-redundant test sets created by removing any remote homologs (TEST2020, CASP12-FM, and CASP13-FM) or by 25% sequence identity cutoff (TEST2018). The large improvement of our method over any single-sequence-based methods for all structural properties is observed for all five test sets (TEST2018, TEST2020, CASP12-FM, and CASP13-FM). More importantly, we demonstrated that an alignment-free method can match or improve over an alignment-based method in predicting 1D structural properties, regardless if it is secondary-structure or tertiary-structure-based 1D properties.

To enlarge our test sets, TEST2020 contains low-resolution structures. To examine if these low-quality proteins affect our conclusion above, we also obtained TEST2020-HQ. As shown in Supplementary Table S5, we found that the performance on TEST2020 is essentially the same as the performance on TEST2020-HQ for all structural properties predicted.

The alignment-free method proposed here can skip the intensive computing time required to search for homologous sequences from an exponentially expanding sequence database. For example, generating PsiBlast sequence profiles and HMM models will require 9.3 hours and 6.9 hours, respectively, for 250 in TEST2018 by utilizing 16 cores of Intel(R) Xeon(R) CPU E5-2620 v4 @ 2.10GHz machine. After that, it takes additional 0.23 hours by NetSurfP-2.0 or 1.1 hours by SPOT-1D to complete the prediction. SPOT-1D, an ensemble of six different models, uses SPOT-Contact, SPIDER3, CCMpred and DCA as input. This makes the pipeline for SPOT-1D extensively time-consuming. By comparison, SPOT-1D-LM takes a total 0.29 hours on the same 16-core CPU for complete prediction with the same or better accuracy. The single-sequence method SPOT-1D-Single is quicker than SPOT-1D-LM (0.04 hours) but with poorer performance. Moreover, SPOT-1D-LM can complete the whole prediction on a Titan X GPU for 0.04 hours only. Thus, it is now feasible for making highly accurate genome-scale prediction on protein secondary and tertiary structural features.

This method is limited to a protein of *≤* 1024 amino-acid residues. This should not hamper the analysis of protein sequence features because the largest structural domain found so far contains 692 amino acid residues^28,29^ with the majority <200 residues. Large (long) proteins usually are made of multiple, mostly independent structural domains connected by intrinsically disorder regions. Thus, it is possible to divide a protein into shorter domains prior to make secondary structure or other structural property prediction by using protein domain prediction tools^30^.

The successful matching performance between alignment-free and alignment-based methods highlights the potential of using a similar combination of language models for other structural properties such as proteins intrinsic disorder^31^ and distance-based contact maps^8,32^ as well as for end-to-end tertiary structure prediction^1,33,34^. In particular, AlphaFold2 has successfully predicted protein structures at an experimental accuracy in CASP14 for those proteins with a minimal of 30 homologous sequences^1^. Our results indicated the possibility that the success of AlphaFold2 can expand to the proteins without homologous sequences by using a combination of language models as input, rather than the multiple aligned homologous sequences as an input.

## Methods

### Datasets

The training and test datasets employed in this work are from our previous work for developing SPOT-1D-Single. Briefly, we started with the dataset prepared by ProteinNet at the highest sequence identity cutoff of 95% according to mmseqs2 tool^35^ to maximize the training data. This leads to 50914 proteins submitted to PDB before the year 2016 with resolution <2.5Å.

To avoid overfitting and achieve an effective validation, we randomly selected 100 proteins one by one from the training set and compared their Hidden Markov Model (HMM) against the HMM of all other proteins in the training set at an e-value cutoff of less than 0.1. Any proteins that were remotely similar to the 100 validation proteins were removed from the training set. In addition, we removed any proteins with length more than 1024. This led to 38913 proteins for training and 99 proteins for validation.

The first test set employed is TEST2018^5^ with 250 proteins released between January 01, 2018 and June 17, 2018 with resolution < 2.5Å and R-free < 0.25, and have sequence similarity less than 25% to all pre-2018 proteins. We further obtained a hard test set TEST2020 that includes all proteins released between May 2018 and April 2020 with removal of close and remote homologs using HMM models to all proteins released before 2018 on PDB. Due to the limitation of the language model used, we further removed the proteins with lengths greater than 1024. The final TEST2020 contains 671 proteins. A further resolution constraint of <2.5Å and R-free<0.25 led to 124 proteins forming TEST2020-HQ.

Apart from the above-mentioned test sets, we also employed independent test sets CASP12-FM (released in year 2016) and CASP13-FM (released in year 2018). These test sets include the free-modelling proteins released during CASP12 and CASP13. CASP12-FM includes 22 proteins and CASP13-FM includes 17 proteins. Free modeling targets are those proteins without known structural templates in the protein databank at the time of releases, which are after all proteins in the training and validation sets.

### Input Features

As shown in Figure 4, we employed the one-hot encoding from the protein sequence concatenated to the language model embeddings generated using ESM-1b and ProtTrans models. The one-hot encoding has a dimension of L *×* 20, where L is the length of the protein. The embedding from ESM-1b is generated from a model trained on the Uniref50 dataset and has a dimension of L *×* 1280. The ProtTrans model was also trained on the Uniref50 and employed the T5-XL model to generate an embedding of dimension of L *×* 1024. Concatenating all these features yielded the final input features of dimension L *×* 2324. This input was utilized for both classification and regression models.

**Figure 4.**
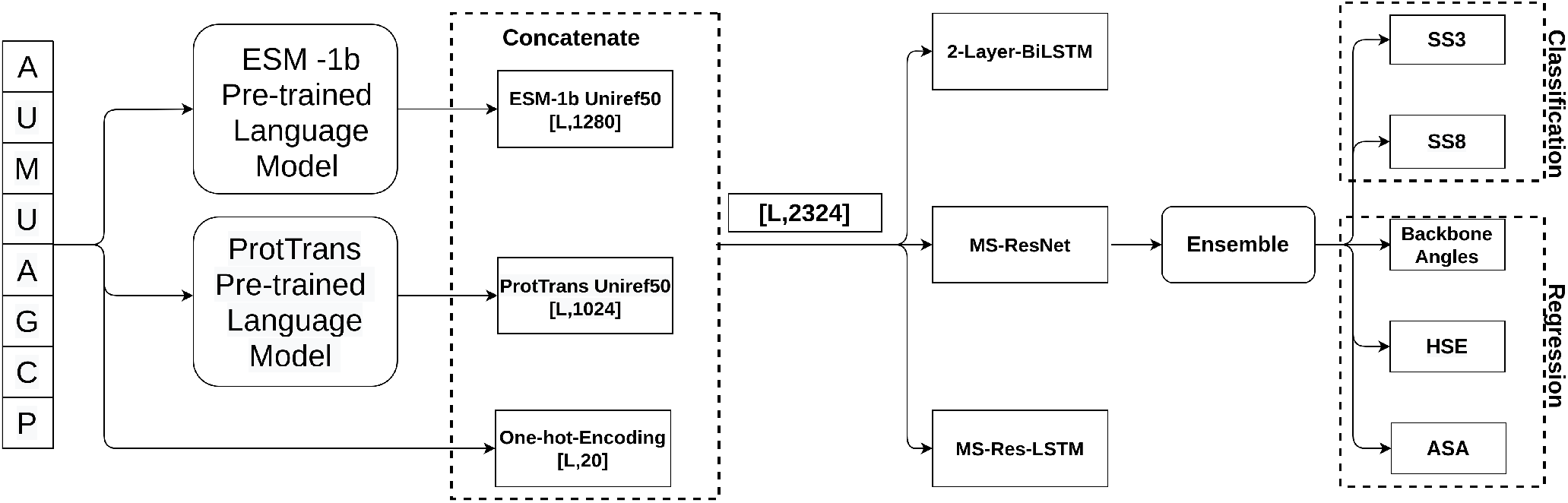
Overview of the model architecture.

### Outputs

The outputs of this method have been divided into two main categories: classification and regression. The classification output is extracted from the classification models with 11 output nodes dedicated for protein secondary structure. We use the Dictionary of Secondary Structure of Proteins (DSSP) for assigning three-state (SS3) and eight-state (SS8) secondary structures^36^. We also predicted 1D structural properties which fall under the regression category. They include the solvent accessible surface area (ASA), protein backbone angles (*ψ, ϕ, θ*, and *τ*), half-sphere exposures (HSE), and contact number (CN). These outputs are identical to the predictions in our previous methods SPOT-1D^5^ and SPOT-1D-Single^14^.

### Neural network architecture

The model utilized in SPOT-1D-LM follows the neural networks utilized in SPOT-1D-Single^14^. In brief, we employed an ensemble of three neural network architectures: 2-layer BiLSTM, multi-scale-ResNet (MS-ResNet), and multi-scale-Res-LSTM (MS-Res-LSTM). The ensemble of LSTM-BRNN and ResNet-based models help the models to identify short- and long-range context throughout the sequence^5^. In total, we trained three models to form an ensemble of three for the classification tasks and regression tasks, respectively. Similar to SPOT-1D-Single, both classification and regression models were trained on a batch size of ten with cross-entropy loss and L1-loss, respectively. The ensemble of classification models employed the mean of the classification probabilities from each model. The mean was also employed for the ensemble of the ASA, HSE-u, HSE-d and CN regression models. For the angle prediction, we utilized the median as in SPOT-1D^5^ to avoid forbidden angle regions.

The first model we trained is a two-layered bidrectional-LSTM with hidden dimension of 1024 followed by two fully connected layers of size 1000^17^. A dropout rate of 0.5 after each LSTM layer was used to avoid overfitting. The second model we trained is a MS-Resnet, which is made of three parallel stacks of ResNet architectures with a great performance for similar tasks^5,8^. The three stacks differ from each other in terms of the kernel size. The first, second, and third stacks of the ResNets have the kernel sizes of three, five, and seven, respectively. Each parallel stack has 15 blocks of ResNet for which the sizes of convolutional layers vary after every five blocks from 64-256. At the end, the output from all three stacks is then concatenated and passed through the output layer. In every ResNet block, we normalized and activated the output of each convolutional layer by applying batch normalization and ReLU activation function^37,38^. We also applied a dropout rate of 0.5 in each block. The third model we trained is MS-Res-LSTM. This model is a hybrid of the first two models. It includes the MS-ResNet in which one parallel stack of three is replaced by four bidirectional-LSTM layers of a hidden size of 128. The ResNet block stacks have the same configuration as the MS-ResNet stacks with kernel sizes of 5 and 7, respectively. A dropout rate of 0.5 was employed in the bidirectional-LSTM layer.

### Performance evaluation

The three-state (SS3) and eight-state (SS8) secondary-structure predictions were evaluated based on the percentage accuracy by concatenating all the proteins together and making an overall assessment. Prediction of ASA, HSE-u, HSE-d, and CN were evaluated by calculating the Pearson’s Correlation Coefficient (PCC) between true and predicted values for each protein and then averaged over the whole dataset^39^. To evaluate the model performance for the backbone angles (*ψ, ϕ, θ*, and *τ*), we calculate the Mean Absolute Error (MAE) between true angles and predicted angles for the whole dataset concatenated together.

### Methods comparison

SPOT-1D-LM developed here was compared with single-sequence-based predictors SPOT-1D-Single, ProteinUnet, SPIDER-Single3, PSIPRED-Single and ASA-Quick. We also compared our method against profile-based methods SPOT-1D, and NetSurfP-2.0. All above-stated methods have stand-alone programs available online at https://github.com/jas-preet/SPOT-1D-Single, https://codeocean.com/capsule/2521196/tree/v1, https://servers.sparks-lab.org/downloads/SPIDER3-Single_np.tgz, http://bioinfadmin.cs.ucl.ac.uk/downloads/psipred/, http://mamiris.com/GENN+ASAquick.tgz, https://sparks-lab.org/downloads/, and https://services.healthtech.dtu.dk/service.php?NetSurfP-2.0, respectively.

## Acknowledgement

This work was supported by the Australian Research Council DP210101875 to K.P and Y.Z. We gratefully acknowledge the use of the High-Performance Computing Cluster Gowonda to complete this research and the aid of the research cloud resources provided by the Queensland Cyber Infrastructure Foundation (QCIF). We also gratefully acknowledge the support of NVIDIA Corporation with the donation of the Titan V GPU used for this research. The support of Shenzhen Science and Technology Program (Grant No. KQTD20170330155106581) and the Major Program of Shenzhen Bay Laboratory S201101001 is also acknowledged.

## Author contribution

JS^*\**^, KP, and JS designed network architectures, JS^*\**^ prepared the data sets and generated input features. JS^*\**^ did deep learning models training, the results analysis, wrote the manuscript, and build a standalone tool and web API. YZ conceived of the study, participated in the initial design, assisted in result analysis, and drafted the whole manuscript. All authors read, contributed to the discussion, and approved the final manuscript.

