## Supplementary Material for "SPOT-1D-LM: Reaching Alignment-profile-based Accuracy in Predicting Protein Secondary and Tertiary Structural Properties without Alignment"

### Supplementary Material: SPOT-1D-LM: Achieving Alignment-profile-based Accuracy in Prediction Protein Secondary and Tertiary Structural Properties without Alignment.

**Jaspreet Singh**[1,*], **Kuldip Paliwal**[1,*], **Jaswinder Singh**[1], **and Yaoqi Zhou**[2,3,4,*]

[1]Signal Processing Laboratory, School of Engineering and Built Environment, Griffith University, Brisbane, QLD 4111, Australia
[2]Institute for Glycomics, Griffith University, Parklands Dr. Southport, QLD 4222, Australia
[3]Institute for Systems and Physical Biology, Shenzhen Bay Laboratory, Shenzhen 518055, China
[4]Peking University Shenzhen Graduate School, Shenzhen 518055, P.R.China

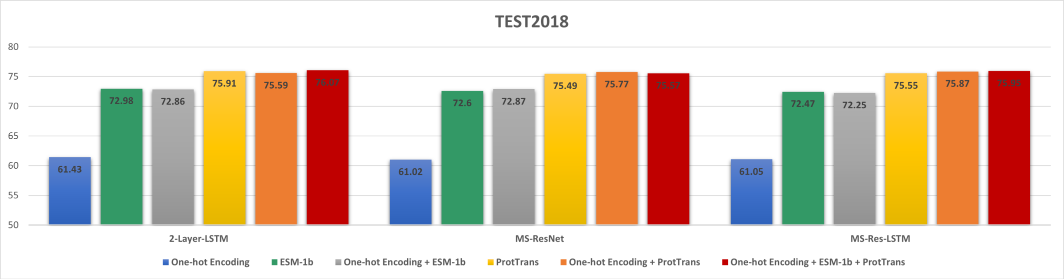


**(a)**

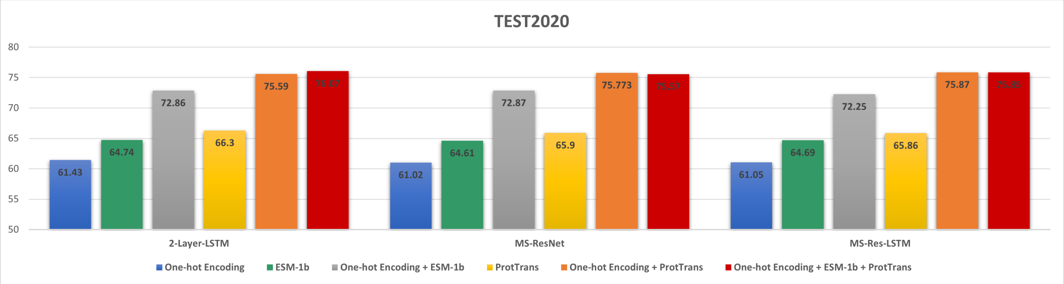


**(b)**

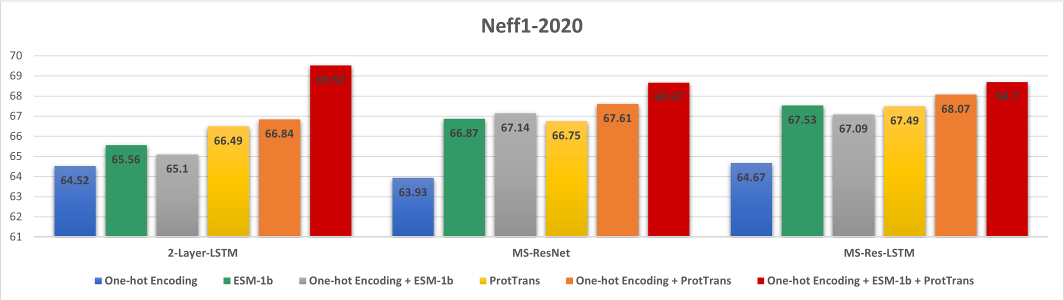


**(c)**

**Supplementary Figure S1:** Performance in eight-state (SS8) secondary structure prediction by using different input features as labelled for three different model architectures on three test sets (TEST2018, TEST2020, and Neff1-2020).

**Supplementary Table S1:** Performance comparison of different architectures based on different input features for ASA, HSE-u, HSE-d, and CN predictions using TEST2018, TEST2020, and Neff1-2020 sets.

| Features | Model | TEST2018 | | | | TEST2020 | | | | Neff1-2020 | | | |
|---|---|---|---|---|---|---|---|---|---|---|---|---|---|
| | | ASA | HSE-U | HSE-D | CN | ASA | HSE-U | HSE-D | CN | ASA | HSE-U | HSE-D | CN |
| One-hot Encoding | 2-Layer-LSTM | 0.668 | 0.547 | 0.549 | 0.573 | 0.637 | 0.387 | 0.466 | 0.482 | 0.653 | 0.368 | 0.502 | 0.519 |
| | MS-ResNet | 0.670 | 0.554 | 0.551 | 0.577 | 0.643 | 0.390 | 0.474 | 0.485 | 0.656 | 0.358 | 0.524 | 0.545 |
| | MS-Res-LSTM | 0.670 | 0.554 | 0.551 | 0.577 | 0.644 | 0.390 | 0.474 | 0.485 | 0.657 | 0.357 | 0.524 | 0.545 |
| One-Hot-Encoding + ESM-1b | 2-Layer-LSTM | 0.788 | 0.717 | 0.722 | 0.766 | 0.702 | 0.475 | 0.558 | 0.578 | 0.694 | 0.427 | 0.588 | 0.603 |
| | MS-ResNet | 0.780 | 0.710 | 0.717 | 0.742 | 0.703 | 0.469 | 0.560 | 0.587 | 0.705 | 0.43 | 0.584 | 0.612 |
| | MS-Res-LSTM | 0.784 | 0.717 | 0.724 | 0.758 | 0.705 | 0.481 | 0.568 | 0.59 | 0.707 | 0.427 | 0.586 | 0.621 |
| One-Hot-Encoding + ProtTrans | 2-Layer-LSTM | 0.808 | 0.744 | 0.76 | 0.785 | 0.723 | 0.512 | 0.592 | 0.609 | 0.723 | 0.469 | 0.614 | 0.628 |
| | MS-ResNet | 0.803 | 0.736 | 0.737 | 0.764 | 0.718 | 0.505 | 0.585 | 0.603 | 0.716 | 0.461 | 0.587 | 0.619 |
| | MS-Res-LSTM | 0.803 | 0.738 | 0.747 | 0.783 | 0.721 | 0.512 | 0.591 | 0.617 | 0.718 | 0.457 | 0.602 | 0.626 |
| ProtTrans + ESM-1b + One-Hot-Encoding | 2-Layer-LSTM | 0.812 | 0.745 | 0.755 | 0.788 | 0.73 | 0.516 | 0.591 | 0.612 | 0.731 | 0.472 | 0.613 | 0.634 |
| | MS-ResNet | 0.807 | 0.748 | 0.748 | 0.786 | 0.722 | 0.51 | 0.588 | 0.607 | 0.719 | 0.444 | 0.599 | 0.622 |
| | MS-Res-LSTM | 0.806 | 0.749 | 0.748 | 0.778 | 0.723 | 0.512 | 0.584 | 0.606 | 0.724 | 0.461 | 0.597 | 0.605 |

**Supplementary Table S2:** Performance comparison of different architectures based on different input features for $\psi$, $\phi$, $\theta$, and $\tau$ predictions on TEST2018, TEST2020, and Neff1-2020 set.

| Features | Model | TEST2018 | | | | TEST2020 | | | | Neff1-2020 | | | |
|---|---|---|---|---|---|---|---|---|---|---|---|---|---|
| | | $\psi$ | $\phi$ | $\theta$ | $\tau$ | $\psi$ | $\phi$ | $\theta$ | $\tau$ | $\psi$ | $\phi$ | $\theta$ | $\tau$ |
| One-hot Encoding | 2-Layer-LSTM | 41.603 | 22.387 | 9.5 | 43.258 | 44.985 | 23.192 | 9.971 | 44.241 | 43.124 | 21.22 | 9.509 | 41.734 |
| | MS-ResNet | 41.307 | 22.509 | 9.532 | 43.382 | 41.307 | 44.202 | 22.766 | 9.904 | 41.548 | 20.717 | 9.286 | 40.657 |
| | MS-Res-LSTM | 41.345 | 22.507 | 9.532 | 43.48 | 44.207 | 22.764 | 9.904 | 43.88 | 41.516 | 20.715 | 9.288 | 40.617 |
| One-Hot-Encoding + ESM-1b | 2-Layer-LSTM | 27.393 | 17.478 | 7.16 | 28.673 | 39.067 | 21.35 | 9.134 | 38.641 | 38.162 | 19.849 | 8.881 | 37.955 |
| | MS-ResNet | 27.737 | 17.675 | 7.247 | 29.139 | 38.316 | 21.281 | 8.964 | 37.881 | 37.832 | 19.704 | 8.726 | 37.338 |
| | MS-Res-LSTM | 27.65 | 17.506 | 7.2 | 28.966 | 38.463 | 21.31 | 8.988 | 38.069 | 37.803 | 19.605 | 8.793 | 37.58 |
| ProtTrans + One-Hot-Encoding | 2-Layer-LSTM | 24.301 | 16.245 | 6.554 | 25.118 | 37.6 | 21.054 | 8.893 | 36.819 | 39.598 | 20.163 | 9.129 | 38.25 |
| | MS-ResNet | 24.477 | 16.396 | 6.613 | 25.336 | 37.281 | 20.945 | 8.781 | 36.443 | 37.933 | 19.859 | 8.753 | 36.494 |
| | MS-Res-LSTM | 24.68 | 16.368 | 6.618 | 25.45 | 37.452 | 20.966 | 8.844 | 36.62 | 37.736 | 19.764 | 8.798 | 36.374 |
| ProtTrans + ESM-1b + One-Hot-Encoding | 2-Layer-LSTM | 23.964 | 16.14 | 6.54 | 24.826 | 36.792 | 20.668 | 8.738 | 36.154 | 36.846 | 19.491 | 8.572 | 36.032 |
| | MS-ResNet | 24.285 | 16.217 | 6.577 | 25.119 | 37.04 | 20.877 | 8.793 | 36.2 | 37.211 | 19.824 | 8.671 | 36.324 |
| | MS-Res-LSTM | 24.396 | 16.428 | 6.617 | 25.285 | 36.877 | 20.846 | 8.725 | 36.124 | 36.746 | 19.42 | 8.569 | 35.488 |

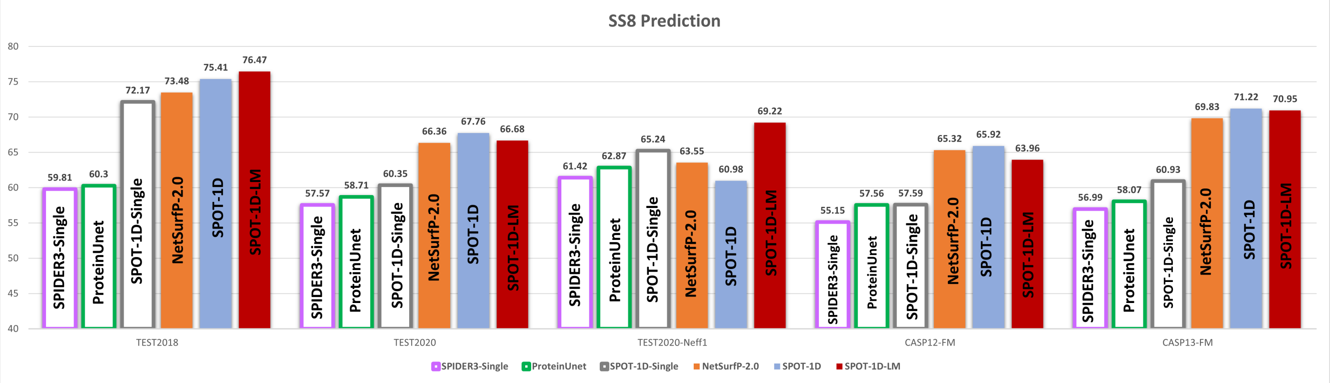


**Supplementary Figure S2:** Comparing the accuracy of eight-state (SS8) secondary structure prediction of SPOT-1D-LM (this work) with single-sequence-based methods (SPIDER3-Single, ProteinUnet, and SPOT-1D-Single) and sequence-profile-based methods (SPOT-1D and NetSurfP-2.0) on five test sets (TEST2018, TEST2020, NEFF1-2020, CASP12-FM, CASP13-FM).

**Supplementary Table S3:** Comparing the accuracy of SPOT-1D-LM (this work) with single-sequence-based methods (SPIDER3-Single, ProteinUnet, and SPOT-1D-Single) and sequence-profile-based methods (SPOT-1D and NetSurfP-2.0) for prediction of secondary structure in three (SS3) and eight (SS8) states, solvent accessibility (ASA), half-sphere-exposure-up (HSE-u), HSE-down (HSE-d), contact number (CN), backbone angles ($\psi$, $\phi$, $\theta$ and $\tau$) for CASP12-FM. Performance measures are accuracy for SS3 and SS8, correlation coefficient for ASA, HSE-u, HSE-d, and CN, and mean absolute errors for the angles.

| Model | SS3 | SS8 | ASA | HSE-u | HSE-d | CN | $\psi$ | $\phi$ | $\theta$ | $\tau$ |
|---|---|---|---|---|---|---|---|---|---|---|
| SPIDER3-Single | 69.15 | 55.15 | 0.586 | 0.516 | 0.467 | 0.554 | 47.462 | 26.168 | 11.639 | 47.591 |
| ProteinUnet | 71.33 | 57.56 | 0.582 | 0.523 | 0.482 | 0.556 | 46.527 | 25.942 | 10.949 | 46.259 |
| SPOT-1D-Single | 72.44 | 57.59 | 0.612 | 0.556 | 0.522 | 0.599 | 43.457 | 25.426 | 10.278 | 44.022 |
| NetSurfP-2.0(profile) | 78.36 | 65.32 | 0.668 | - | - | - | 35.127 | 22.262 | - | - |
| SPOT-1D (profile) | 79.53 | 65.92 | 0.667 | 0.660 | 0.621 | 0.692 | 33.962 | 21.844 | 8.700 | 33.114 |
| SPOT-1D-LM (This work) | 77.45 | 63.96 | 0.674 | 0.661 | 0.629 | 0.686 | 35.955 | 22.29 | 8.87 | 35.236 |

**Supplementary Table S4:** Comparing the accuracy of SPOT-1D-LM (this work) with single-sequence-based methods (SPIDER3-Single, ProteinUnet, and SPOT-1D-Single) and sequence-profile-based methods (SPOT-1D and NetSurfP-2.0) for prediction of secondary structure in three (SS3) and eight (SS8) states, solvent accessibility (ASA), half-sphere-exposure-up (HSE-u), HSE-down (HSE-d), contact number (CN), backbone angles ($\psi$, $\phi$, $\theta$ and $\tau$) for CASP13-FM. Performance measures are accuracy for SS3 and SS8, correlation coefficient for ASA, HSE-u, HSE-d, and CN, and mean absolute errors for the angles.

| Model | SS3 | SS8 | ASA | HSE-u | HSE-d | CN | $\psi$ | $\phi$ | $\theta$ | $\tau$ |
|---|---|---|---|---|---|---|---|---|---|---|
| SPIDER3-Single | 71.26 | 56.99 | 0.565 | 0.462 | 0.408 | 0.495 | 46.164 | 25.315 | 11.076 | 46.348 |
| ProteinUnet | 70.63 | 58.07 | 0.571 | 0.480 | 0.433 | 0.513 | 46.884 | 25.036 | 10.380 | 46.093 |
| SPOT-1D-Single | 73.21 | 60.93 | 0.572 | 0.489 | 0.464 | 0.531 | 45.231 | 25.124 | 9.889 | 44.903 |
| NetSurfP-2.0(profile) | 82.11 | 69.83 | 0.698 | - | - | - | 31.817 | 21.577 | - | - |
| SPOT-1D (profile) | 83.55 | 71.22 | 0.701 | 0.683 | 0.632 | 0.704 | 28.489 | 20.238 | 7.551 | 27.867 |
| SPOT-1D-LM (This work) | 82.154 | 70.946 | 0.763 | 0.753 | 0.711 | 0.785 | 30.68 | 20.092 | 7.563 | 29.634 |

**Supplementary Table S5:** Comparing the accuracy of SPOT-1D-LM (this work) with single-sequence-based methods (SPIDER3-Single, ProteinUnet, and SPOT-1D-Single) and sequence-profile-based methods (SPOT-1D and NetSurfP-2.0) secondary structure in three (SS3) and eight (SS8) states, solvent accessibility (ASA), half-sphere-exposure-up (HSE-u), half-sphere-exposure-down (HSE-d), contact number (CN), backbone angles ($\psi$, $\phi$, $\theta$, and $\tau$) for TEST2020-HQ. Performance measures are accuracy for SS3 and SS8, correlation coefficient for ASA, HSE-u, HSE-d, and CN, and mean absolute errors for the angles.

| Model | SS3 | SS8 | ASA | HSE-u | HSE-d | CN | $\psi$ | $\phi$ | $\theta$ | $\tau$ |
|---|---|---|---|---|---|---|---|---|---|---|
| SPIDER3-Single | 71.02 | 58.23 | 0.612 | 0.451 | 0.469 | 0.504 | 44.051 | 24.264 | 13.061 | 47.424 |
| ProteinUnet | 71.28 | 58.78 | 0.573 | 0.459 | 0.477 | 0.499 | 43.282 | 23.699 | 12.120 | 46.484 |
| SPOT-1D-Single | 72.22 | 59.70 | 0.627 | 0.491 | 0.530 | 0.540 | 42.624 | 23.011 | 11.466 | 45.393 |
| NetSurfP-.2.0 (Profile) | 80.69 | 68.34 | 0.716 | - | - | - | 31.298 | 19.904 | - | - |
| SPOT-1D (profile) | 81.97 | 70.41 | 0.720 | 0.625 | 0.679 | 0.706 | 28.875 | 18.782 | 9.512 | 32.311 |
| SPOT-1D-LM (This work) | 79.70 | 67.73 | 0.755 | 0.624 | 0.666 | 0.704 | 32.011 | 19.523 | 9.743 | 35.271 |
